# Downregulation of the Ca^2+^ sensor Synaptotagmin-1 (*SYT1*) in Parkinson’s Disease: Insights from Gene Expression Profiling

**DOI:** 10.1101/2025.09.12.675906

**Authors:** Giovanna C. Cavalcante, Giordano B. Soares-Souza

## Abstract

Parkinson’s disease (PD) is a neurodegenerative disease characterized by the progressive loss of dopaminergic neurons and by the intracellular accumulation of alpha-synuclein, leading to motor and non-motor symptoms. Despite being a widely studied disease, so new mechanisms should be investigated as possible paths for future diagnostics and treatments in PD. Here, we performed an *in silico* analysis of the global gene expression of tissues from different brain regions (prefrontal area, putamen, and substantia nigra) in PD patients and controls, to demonstrate differentially expressed genes (DEGs). We analyzed the dataset series GSE20295 from GEO, which comprises GSE20168, GSE20291, and GSE20292. We identified 13 DEGs, all exhibiting downregulation in PD tissues compared to controls. Notably, the *SYT1* (Synaptotagmin-1) gene demonstrated the lowest expression level and nearly the most significant adjusted p-value. SYT1 is implicated in calcium ion sensor activity, a functional domain showing substantial fold enrichment in our study. This gene encodes a protein pivotal for neurotransmitter release at synapses. We also found a significant role of calcium-related processes in PD pathology through GSEA, indicating an overall increase in their activity and a disruption in neurotransmitter release mechanisms mediated by Ca^2+^. Despite limited research on the correlation between SYT1 and PD to date, our findings suggest the *SYT1* gene holds promise as a possible target for PD. Further investigations are needed to elucidate this association fully.

## Introduction

Parkinson’s disease (PD) is a neurodegenerative disease characterized by the progressive loss of dopaminergic neurons, especially in the Substantia Nigra (SN) region of the brain, and by the intracellular accumulation of alpha-synuclein, leading to symptoms such as bradykinesia, tremors, postural instability and muscle rigidity, in addition no non-motor symptoms that include cognitive deficit and anxiety (Reich & Savitt, 2019; Jankovic & Tan, 2020; Simon et al., 2020). Considered the second most common neurodegenerative disease in the world, a significant increase in the population affected by PD over 65 years of age is estimated from 1990 to 2019 globally, with an even higher percentage of affected individuals over 80 years of age (Ou et al., 2021).

Despite being a widely studied disease, there are still no markers capable of indicating the early onset of idiopathic PD (Jankovic & Tan, 2020). Therefore, it is imperative that new mechanisms be investigated as possible paths for future diagnostics and treatments in PD. One way to do this is to explore the global expression of genes and verify possible relationships with the development and progression of the disease. Therefore, in the present work we performed an *in silico* analysis of the global gene expression of tissues from different brain regions in PD patients and controls, to demonstrate differentially expressed genes (DEGs).

## Methods

### Sampling

In this study, we analyzed the Gene Expression Omnibus (GEO) dataset series GSE20295, which comprises three datasets of tissues from different brain regions of PD patients (PD) and Controls (CT): GSE20168 (prefrontal area 9, n=29), GSE20291 (putamen, n=35), and GSE20292 (whole substantia nigra, n=29). Using the Affymetrix Human Genome U133A Array platform, the transcriptional analysis of these 93 samples (40 PD patients and 53 controls) was originally performed by Zhang et al. (2005).

### Data analysis

All data analysis was performed in R (R Core Team, 2021) using R Studio (RStudio Team, 2020). For extracting the dataset series GSE20295 from GEO, we used *GEOquery* (Davis and Meltzer, 2007). Microarray analysis was primarily performed using *limma* (Ritchie et al., 2015). For better visualizing the distribution of the samples by different classes, we used the UMAP (Uniform Manifold Approximation and Projection) plot generated with *umap* (McInnes et al., 2018). Volcano plot with the adjustment of the p-value with false discovery rate (FDR) method and the log2foldchange (logFC) was plotted using *ggplot2* (Wickham, 2016) and *EnhancedVolcano* (Blighe et al., 2023).

We then employed the following public online platforms for further assessment: GTEx (www.gtexportal.org), STRING (www.string-db.org), The Human Protein Atlas (www.proteinatlas.org) and Gene4PD (www.genemed.tech/gene4pd). In addition, Gene Ontology (GO) enrichment analysis for biological processes (BP) was performed with *clusterProfiler* (Yu et al., 2012; Wu et al., 2021) and, for molecular functions (MF), we used ShinyGO v.0.80 (Ge et al., 2020). To identify terms and pathways with consistent under- or overexpression, we conducted the gene-set enrichment analysis (GSEA) using *clusterProfiler* for GO and KEGG (Kyoto Encyclopedia of Genes and Genomes) databases.

## Results and Discussion

### Distribution analysis

Here, we investigated the differential gene expression (DGE) between case and control, considering tissues from three different brain regions: prefrontal cortex, putamen, and substantia nigra (SN). UMAP was performed to better visualize the sample distribution across these three regions (Figure 1). The UMAP visualization effectively captures the distinct spatial organization of samples from three different brain regions, and the clear separation observed in the UMAP plot reflects the inherent differences in gene expression or molecular profiles among these regions, particularly between SN and the other two regions.

**Figure 1.**
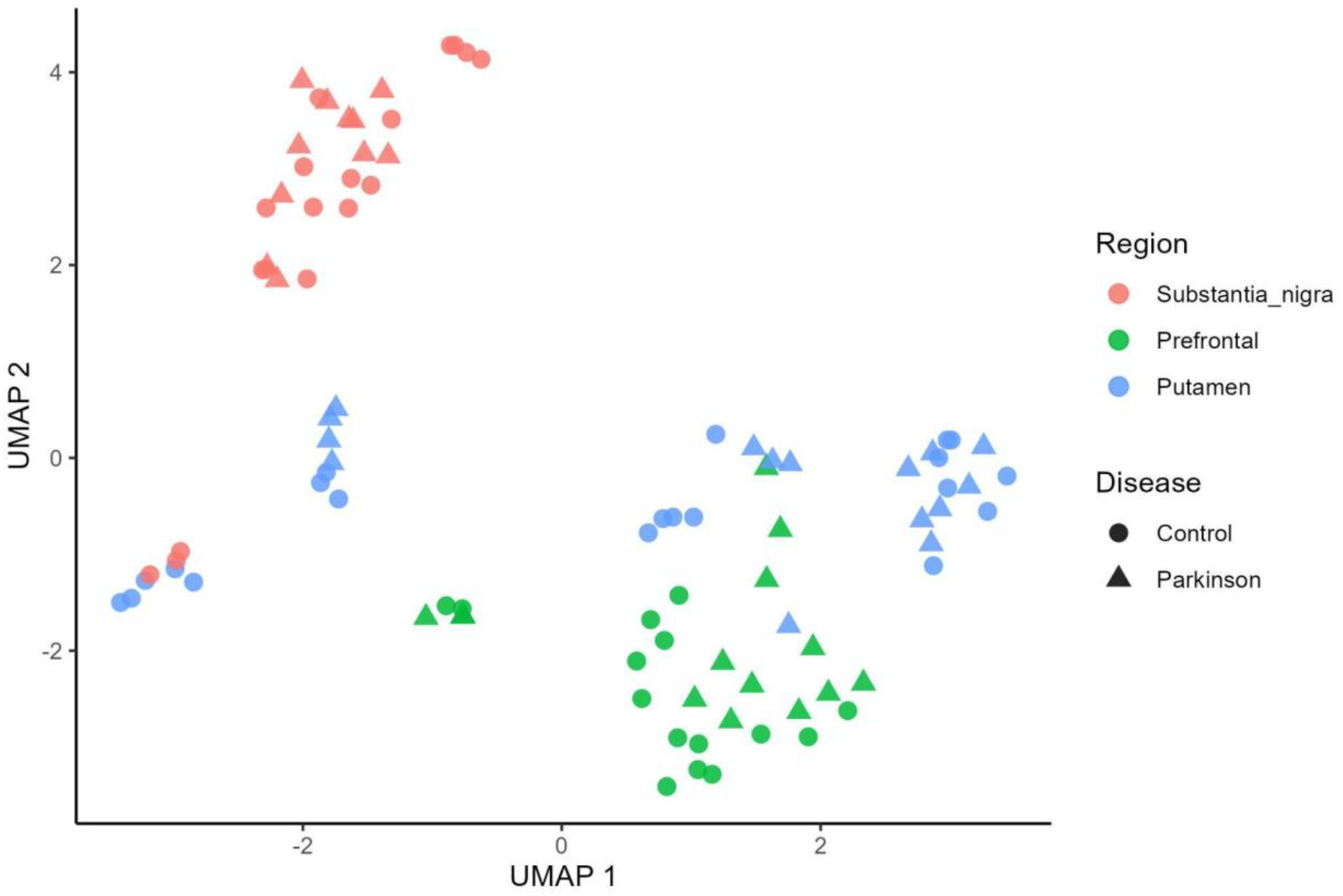
UMAP for the spatial distribution of samples from the three brain regions for case and control.

### DGE analysis

We systematically evaluated various combinations of case-control comparisons across the three distinct brain regions, aiming to comprehensively assess the differential molecular signatures associated with each region. As expected from the UMAP, individual comparisons between brain regions yielded DEGs (e.g., putamen vs substantia nigra in the case group), reaffirming unique molecular profiles for each region. However, no DEGs were identified in comparisons between the same tissues in the case and control groups (e.g., prefrontal in the case vs. control groups). Therefore, we opted to proceed only with the case-control comparison encompassing all three brain regions within each group, providing an assessment of molecular distinctions across the entire dataset (Figure 2).

**Figure 2.**
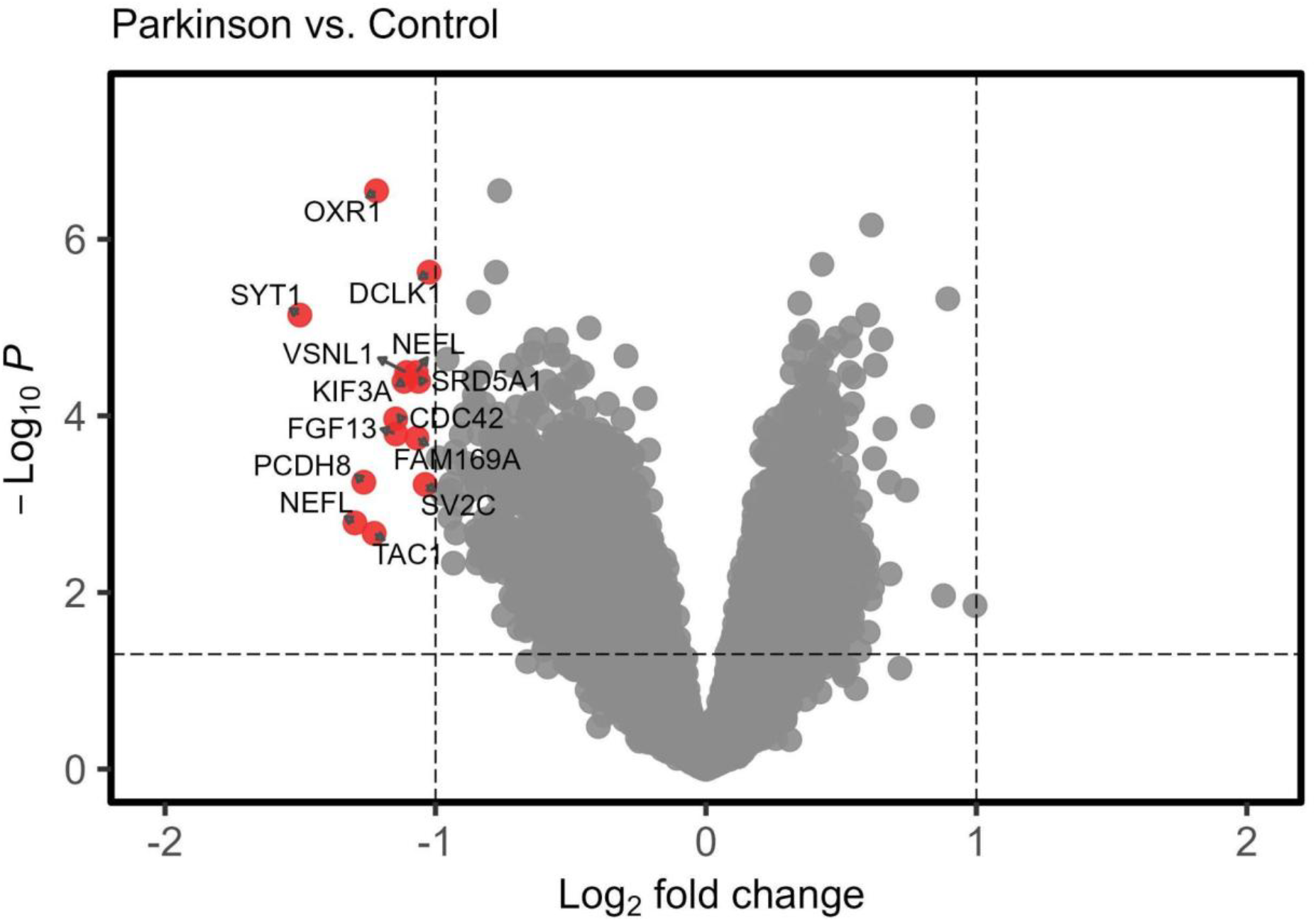
Volcano plot of the gene expression (p<0,05 with FDR adjustment and logFC=1) for the case and control groups.

We found 14 probes representing genes that were DE, but two of them target the same gene, *NEFL*. Based on the adjusted p-value, we considered the most significant one (221801_x_at). Regardless, Table 1 presents all found DEGs and their respective microarray probes, logFC, p-value and adjusted p-value by FDR. All of them were downregulated in the PD tissues in comparison to the CT tissues. Interestingly, *SYT1* (Synaptotagmin-1) gene presented the lowest expression and nearly the most significant adjusted p-value.

**Table 1.**
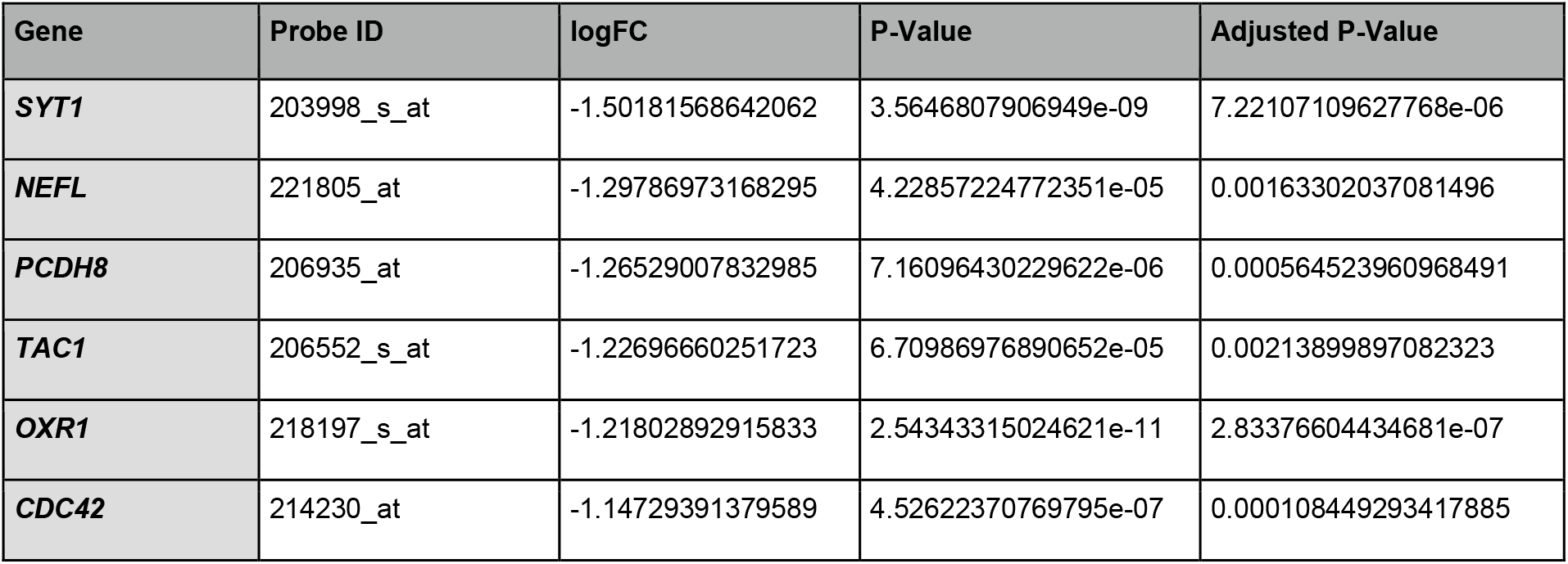
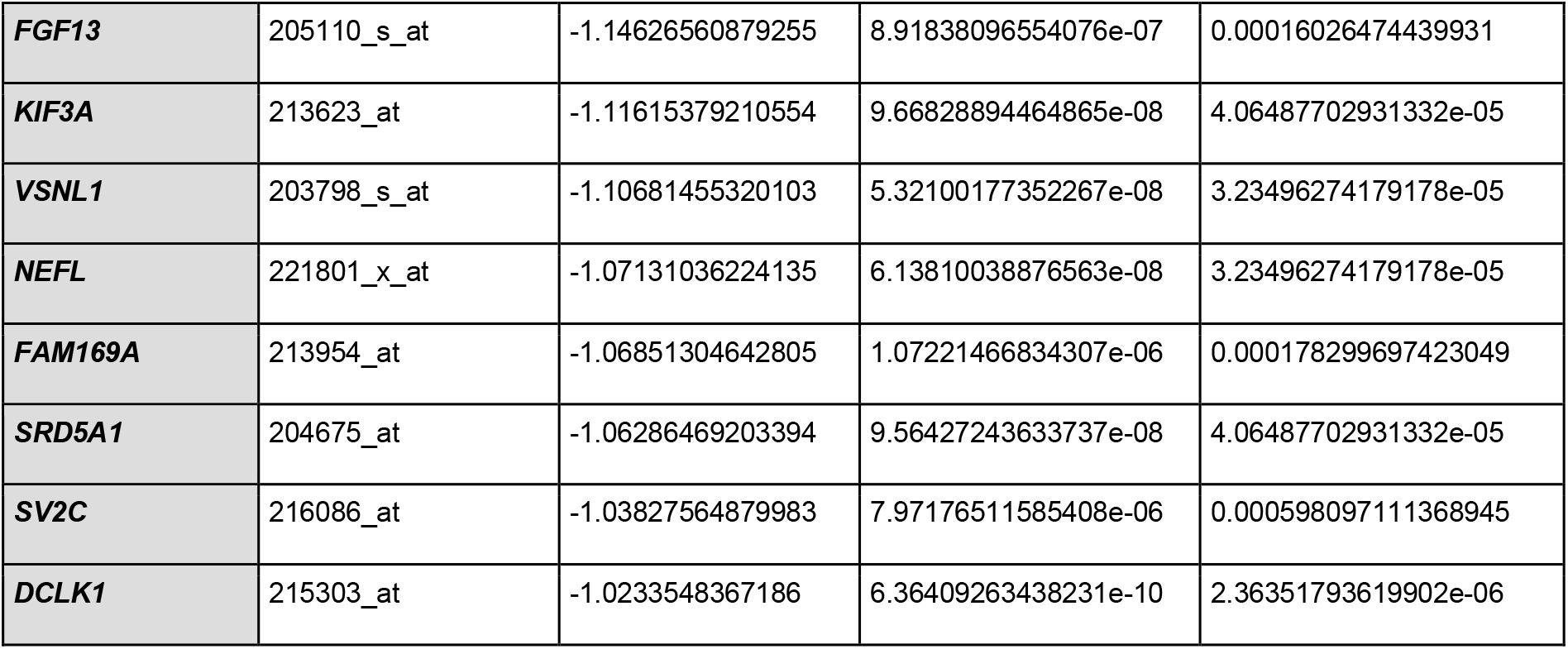
Characterization of the differences in gene expression between case and control.

The *SYT1* gene encodes a homonymous protein that is normally highly expressed in the brain, as seen in Figure 3A in comparison to whole blood, due to its crucial role in neurotransmitter release at synapse, as demonstrated by some of the proteins associated with SYT1 in Figure 3B. This is reinforced by the SYT1 protein expression in different cell types, mostly neurons and glial cells (Figure 3C). Importantly, SYT1 triggers synaptic vesicle fusion upon calcium cellular influx, which is essential for synaptic transmission (Fernández-Chacón et al., 2001; Südhof, 2013).

**Figure 3.**
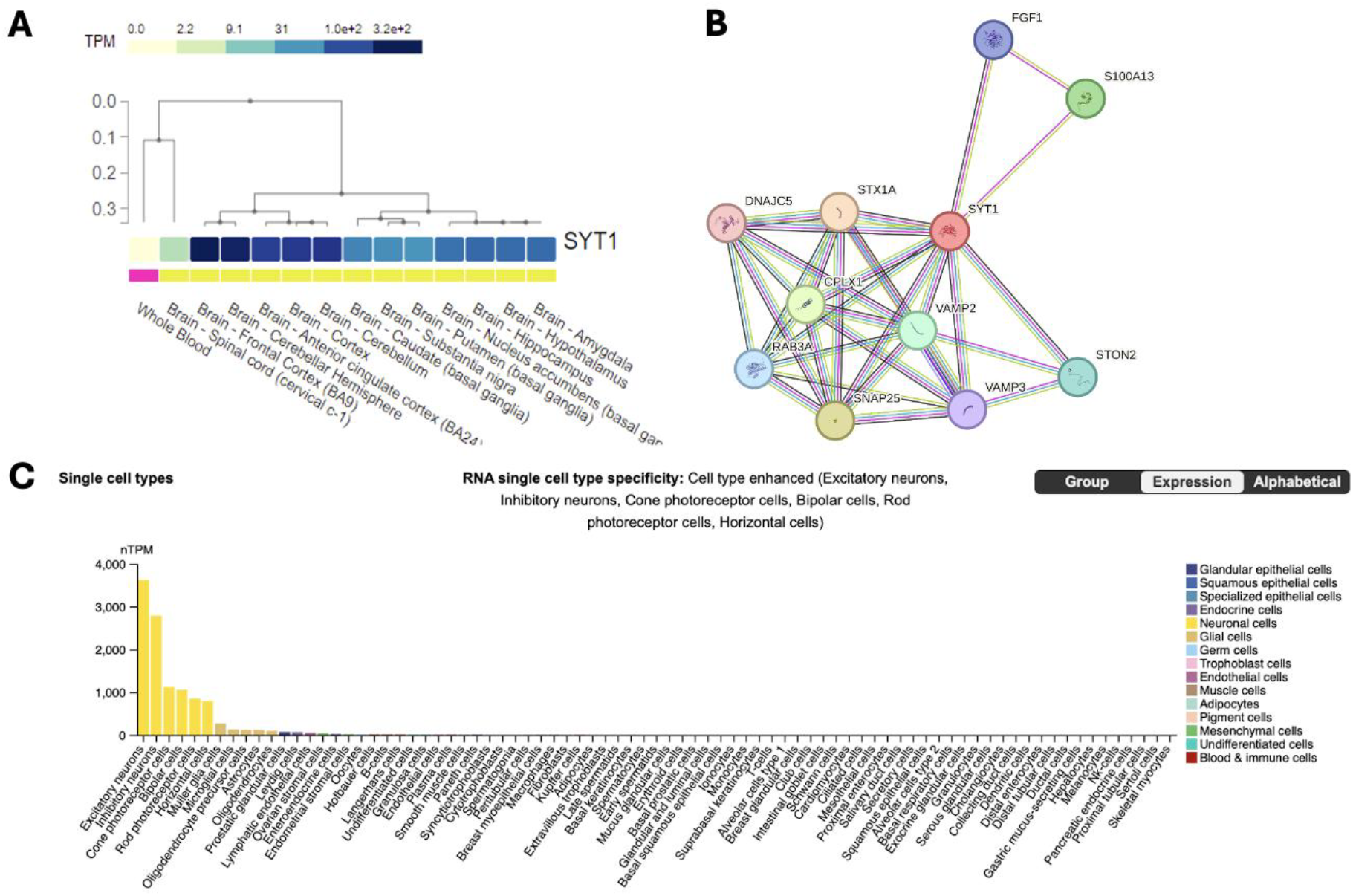
**A)** Expression of *SYT1* gene in 13 different brain tissues, as well as whole blood, from healthy individuals at GTEx Portal (www.gtexportal.org). **B)** Network of proteins associated with SYT1 generated by STRING (www.string-db.org). **C)** RNA single cell type specificity for SYT1 from The Human Protein Atlas (www.proteinatlas.org).

Our observation aligns with a prior investigation conducted with microarray analysis, sourced from the Gene4PD platform (www.genemed.tech/gene4pd). That study also demonstrated the downregulation of *SYT1* gene in PD compared to controls in a different cohort, along with findings such as calcium signaling and mitochondrial pathways (Dumitriu et al., 2016). Interestingly, a work measured SYT1 levels in cerebrospinal fluid (CSF) from 135 PD patients and found it to be decreased in comparison to controls (Brinkmalm et al., 2020). These other SYT1 results at least strengthen its significance in the disease’s pathogenesis.

### Enrichment analysis

Then, we performed the GO enrichment analysis for biological processes of the significant DEGs (Figure 4). Most of the processes found are involved, as expected, with synapse pathways and development of different areas of the brain, as well as pathways involved with transport along microtubules.

**Figure 4.**
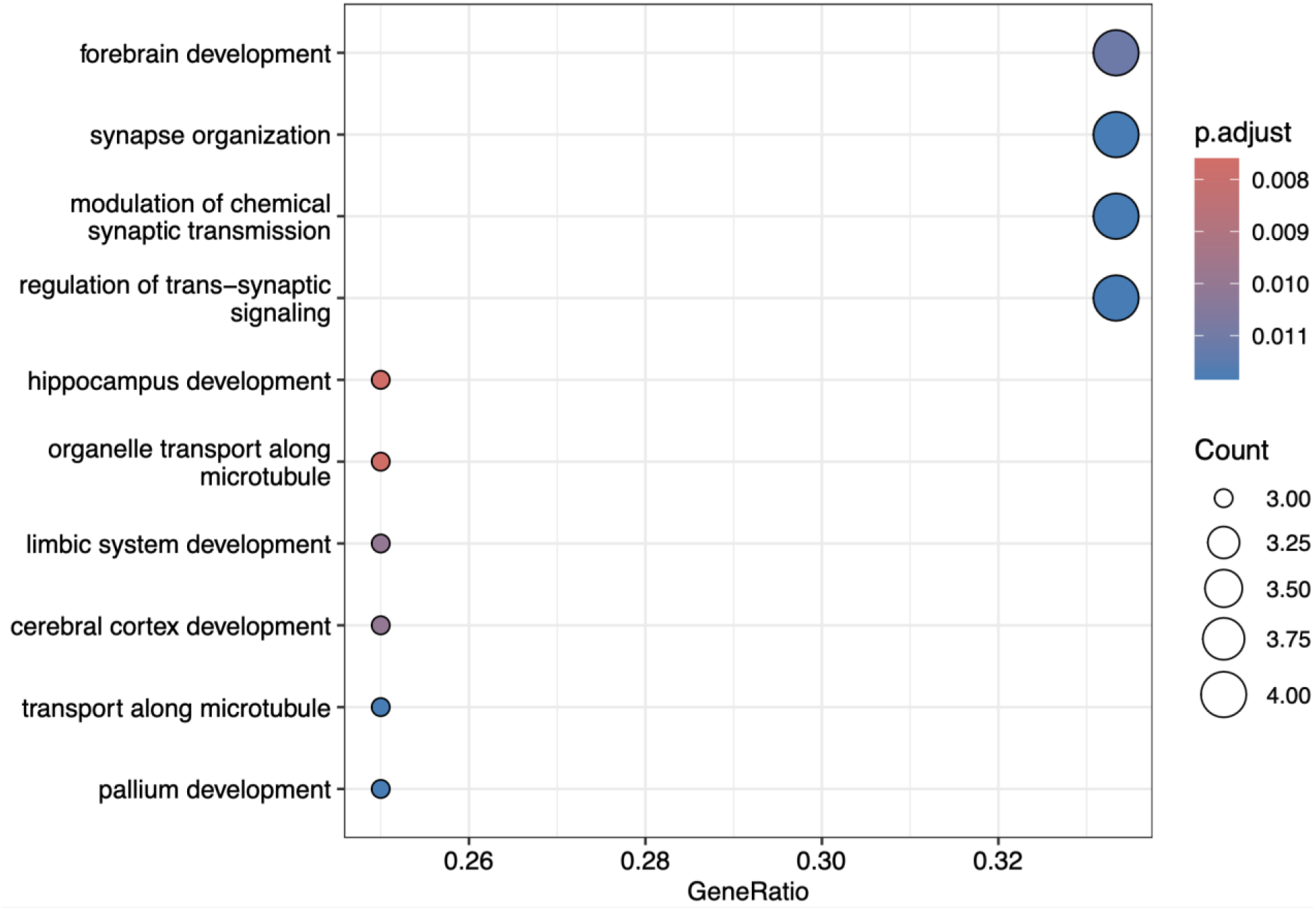
GO enrichment analysis for biological processes of the 13 found DEGs.

In the examination of the GO molecular function of the genes listed in Table 2, we highlight the involvement of three genes, namely *KIF3A, FGF13*, and *DCLK1*, in processes associated with microtubule function. Additionally, we observe that *SYT1*, the gene exhibiting the most pronounced downregulation in our analysis, is implicated in a functional domain with substantial fold enrichment: calcium ion sensor activity.

**Table 2.** GO Molecular function of the 13 DEGs found in our study.

| Enrichment FDR | Number of Genes | Pathway Genes | Fold Enrichment | Pathway | Genes |
| --- | --- | --- | --- | --- | --- |
| 0.0317917685538915 | 1 | 3 | 586.692307692308 | GO:0099184 - structural constituent of postsynaptic intermediate filament cytoskeleton | *NEFL* |
| 0.0317917685538915 | 1 | 4 | 440.019230769231 | GO:0003865 - 3-oxo-5-alpha-steroid 4-dehydrogenase activity | *SRD5A1* |
| 0.0317917685538915 | 1 | 4 | 440.019230769231 | GO:0033765 - steroid dehydrogenase activity acting on the CH-CH group of donors | *SRD5A1* |
| 0.0317917685538915 | 1 | 4 | 440.019230769231 | GO:0047751 - cholestenone 5-alpha-reductase activity | *SRD5A1* |
| 0.0317917685538915 | 1 | 4 | 440.019230769231 | GO:0061891 - calcium ion sensor activity | *SYT1* |
| 0.0417048189312471 | 1 | 6 | 293.346153846154 | GO:0035671 - enone reductase activity | *SRD5A1* |
| 0.0317917685538915 | 3 | 286 | 18.4623453469607 | GO:0008017 - microtubule binding | *KIF3A*,<br>*FGF13*,<br>*DCLK1* |
| 0.0317917685538915 | 3 | 400 | 13.2005769230769 | GO:0015631 - tubulin binding | *KIF3A*,<br>*FGF13*,<br>*DCLK1* |

SYT1 is an essential Ca^2+^ sensor in both synchronous and asynchronous neurotransmitter release in synapse, presenting two Ca^2+^-binding C2 domains (C_2_A and C_2_B) (Bowers et al., 2020; Shields et al., 2020) and an exocytosis/endocytosis bidirectional function in synaptic transmission (Chen et al., 2022). Indeed, through the execution of GSEA, we discerned that 13 GO biological processes associated with calcium exhibited enrichment, predominantly upregulated, but we observed the downregulation of calcium ion-regulated exocytosis of neurotransmitter (enrichment score = -0.61; adjusted p-value = 0.041). Furthermore, our GSEA focusing on KEGG revealed enrichment in the calcium signaling pathway (enrichment score = 0.40; adjusted p-value = 2.408e-08).

These findings indicate a significant role of calcium-related processes in PD pathology, implying an overall increase in their activity, but suggesting a disruption in neurotransmitter release mechanisms mediated by Ca^2+^. In this context, despite the longstanding recognition of SYT1 as a calcium ion sensor, it is clear that ongoing research continues to unveil novel aspects of its function in Ca^2+^-mediated signaling pathways, as excessive calcium in the brain has been related to PD (Lautenschläger et al., 2018).

To address this idea, considering mitochondria are known to play key roles in PD, we also assessed the gene expression of mitochondrial calcium influx and efflux channels. Although not statistically significant considering the adjusted p-value, we found reduced expression of two mitochondrial calcium influx genes (*MICU1* and *MICU2*) and increased expression of the mitochondrial calcium efflux gene *SLC8B1* (which encodes NCLX) in PD brain tissues compared to controls (Supplementary Table S1). The parallel downregulation of *SYT1* reinforces disrupted calcium signaling, potentially impairing synaptic transmission in PD. Together, these observations might hint at a broader calcium dysregulation mechanism in PD, affecting both mitochondrial function and neurotransmitter release.

### SYT1 regulation and Parkinson’s Disease

To date, limited research has investigated the correlation between SYT1 and PD. For instance, a set of four proteins including SYT1 has also been suggested as potential biomarker for PD due to an association with immune infiltration and regulation by the microRNA-92a family (Zhang et al., 2023), which in turn has been previously related to PD (Taguchi and Wang, 2018). Besides miR-92a, it is noteworthy that miR-34a has been associated with PD (Grossi et al., 2021), and that this miRNA has also been reported targeting *SYT1* in neurogenesis (Morgado et al., 2015).

Recently, it has been shown that conditional deletion of SYT1 in dopamine neurons does not impair some dopamine-dependent motor tasks in mice, suggesting that these behaviors can be sustained by basal dopamine levels (Delignat-Lavaud et al., 2023). On the other hand, another recent study has shown SYT1 downregulation in early synucleinopathy in rats, as evidenced by decreased expression of the *SYT1* transcript in neurons containing phosphorylated α-synuclein (pSyn) inclusions (Patterson et al., 2024). This suggests dysfunction in synaptic vesicle release machinery within affected neurons and that the downregulation of SYT1 in early synucleinopathy highlights its potential role in the pathogenesis of PD.

## Conclusions

In this study, we identified the downregulation of *SYT1* in Parkinson’s disease (PD) cases compared to controls, suggesting its potential as a future biomarker in PD pathogenesis due to its role in synaptic transmission. Additionally, we observed decreased expression of calcium ion-regulated exocytosis of neurotransmitter processes, implicating *SYT1* involvement. Limitations include the potential reflection of synaptic loss rather than causative factors due to the absence of neuronal/synaptic quantification, and the non-specificity of *SYT1* expression alteration to PD. Finally, we recommend further research with larger cohorts or more sensitive methods to enhance understanding of *SYT1* in PD pathophysiology.

## Author contributions

GCC: Conceptualization; Data curation; Formal analysis; Investigation; Methodology; Project administration; Writing - original draft; and Writing - review & editing. GBSS: Data curation; Formal analysis; Investigation; Methodology; Visualization; and Writing - review & editing

## Acknowledgements

The authors thank Dr. Alicia J. Kowaltowski (University of São Paulo, Brazil) and Dr. Patricia M. de Carvalho Aguiar (Albert Einstein Israelite Hospital, Brazil) for their insights in the process.

